# Aperture Phase Modulation with Adaptive Optics: A Novel Approach for Speckle Reduction and Structure Extraction in Optical Coherence Tomography

**DOI:** 10.1101/406108

**Authors:** Pengfei Zhang, Suman K. Manna, Eric B. Miller, Yifan Jian, Ratheesh Kumar Meleppat, Marinko V. Sarunic, Edward N. Pugh, Robert J. Zawadzki

## Abstract

Speckle is an inevitable consequence of the use of coherent light in optical coherence tomography (OCT), and often acts as noise that obscures micro-structures of biological tissue. We here present a novel method of suppressing speckle noise intrinsically compatible with adaptive optics (AO) in OCT system: by modulating the phase inside the imaging system pupil aperture with a segmented deformable mirror, thus producing minor perturbations in the point spread function (PSF) to create un-correlated speckle pattern between B-scans, and further averaging to wash out the speckle but maintain the structures. It is a well-controlled and universal method which can efficiently determine the optimal range of phase modulation that minimizing speckle noise while maximizing image resolution and signal strength for different systems and/or samples. As an active method, its effectiveness and efficiency were demonstrated by both ex-vivo non-biological and in-vivo biological applications.

## Introduction

Speckle is a long-standing issue in all imaging technologies that use coherent light sources^1-4^, arised from the interference between light scattered by a random distributed scatters inside the system point-spread function (PSF), and observed as voxel-to-voxel intensity fluctuations in the image^5,6^. Although speckle is a potential useful information about the dynamics of sample microstructure, in most applications it acts as a major noise source that degrades image quality. Optical coherence tomography (OCT) is a volumetric imaging technology developed in 1991^7^. It has been soon adopted in many biomedical applications^8-12^. However, as a method dependent on the coherent properties of light, OCT images suffer from speckle noise^13-15^.

Many approaches have been taken to suppress speckle, including generation of multiple images by various means with uncorrelated speckle patterns, followed by averaging^16-19^. A weakness of these methods is that the number of uncorrelated speckle patterns that can be created is typically small, limiting speckle suppression by averaging. Speckle reduction methods using digital post-processing have also been proposed^20-23^. However, digital post-processing usually reduces speckle by spatial averaging or filtering, which necessarily reduces image resolution. Recently, it was shown that simple averaging of suitably numerous, well aligned images can reduce speckle for *in vivo* imaging, and it was hypothesized that subcellular motility of scatters was responsible for varying the speckle pattern between frames^24,25^. Because this latter method relies on time-dependent variation in the sample microstructure, it is inherently passive and dependent on the underlying time course of the mobile scatters. As a way of potentially overcoming the limitations of passive averaging, speckle modulating OCT (SM-OCT) was recently developed^26^. By introduction of a ground-glass diffuser in the external optical path, the method generates random, time-varying changes in the sample beam. The authors hypothesize that their approach introduces axial phase variation in the imaging plane, but this is not directly under experimenter control, and the phase shift cannot be readily repeated. In contrast, as characterized in classical optical theory and applied in adaptive optics (AO) imaging^27^, the phase can be precisely controlled by manipulation of the wavefront corrector at the system pupil aperture, and this suggests the possibility of using AO technology to create a method for speckle suppression that would be readily controllable and broadly applicable to OCT.

The core of all AO-enhanced imaging is the active control of the wavefront at the system aperture, a controlled implementation mostly by means of a deformable mirror (DM) or spatial light modulators (SLMs) to optimize the wavefront over the pupil to allow the system to operate at diffraction-limited performance^28-31^. Here, we take advantage of this exquisite control to create a novel method for speckle noise reduction aperture phase modulation AO-OCT (APM-AOOCT). This method employs sub-micron piston modulations of the DM segments to introduce random phase variation for all segments in both spatial and temporal directions. In describing APM-AO-OCT, we first present the hypothesized underlying mechanism, namely that the modulations of DM mirror segments about their AO-optimized positions slightly alter the PSF, randomizing over samples the contributions from different scatters to create uncorrelated speckle pattern, so that averaging can efficiently reduce the speckle. We then address the inherent conflict between speckle noise reduction and preservation of signal resolution and strength by determining an optimum mirror segment displacement range. We further identify a relatively small subset of the total set of mirror configurations within this range that maximally reduce speckle while preserve resolution and signal strength. Finally, we demonstrate the success of APM-AO-OCT with *in vivo* mouse retina imaging applications.

## Methods

### (1)AO-OCT system configuration

In adaptive optics (AO) systems used in vision science^27^, a deformable mirror (DM) is placed in an optical plane conjugate with the pupil aperture to correct aberrations of the cornea and lens, as shown in Fig. 1 (a), which provides a schematic of the sample arm of our AO-OCT system built for *in vivo* mouse retinal imaging. The DM (PTT111, IRIS AO, Inc., photo in Fig. 1 (a), bottom right inset) has 37 segments with 111 actuators (3 actuators per segment to independently control the displacement/piston, tip and tilt). The DM segments have nanometer level displacement resolution (z-offset of the mirror surface) with a working range of [-2, 2]μ (Supplementary information (SI): Fig_S1). When the DM operates in ‘flat’ mode, the displacements of all segments are zero (Fig. 1 (b)). The piston of each segment can be independently controlled to operate in a ‘random’ mode (Fig. 1 (c)).

**Fig. 1.**
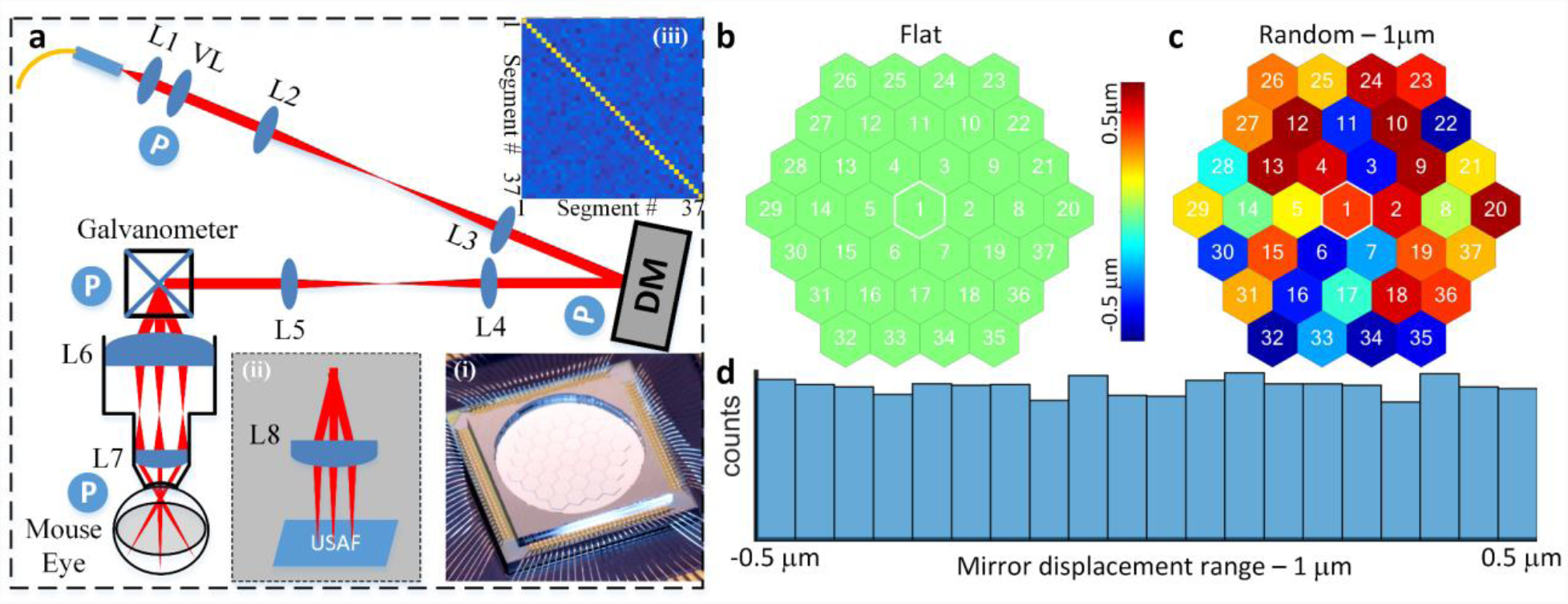
System setup (photo in SI: Fig. S2) **and the geometry of the segmented deformable mirror**. **a**, OCT sample arm (Inset: (**i**) Photo of the DM; (**ii**) setup for USAF 1951 resolution test target; (**iii**) covariance analysis of the random segments pistons). **b**, Mirror configuration – Flat. **c**, Mirror Configuration mirror segments with random displacements [SI: Video_S1]. **d**, Histogram of the mirror displacements for 100 mirror configurations in which the displacement range was 1mm (0 ± 0.5 μ m); Abbreviations: L#: Lens, VL: variable focus length liquid lens, DM: Deformable mirror, USAF: 1951 USAF (U.S. Airforce) resolution test target, P (circled in blue) optical planes conjugate with the pupil.

The lenses used in the sample arm are VIS-NIR coated achromatic lenses (400-1000 nm, Edmunds Optics, key parameters are shown in SI: Table S1). A reference arm was built with dispersion compensation prisms. A SLD (T870-HP, Superlum, ranged from [780, 960] nm and centered at 870 nm) served as the light source for NIR OCT with a power at mouse pupil of 900 mW; A customized spectrometer with 2048 pixels was used to acquire the OCT spectra.

### (2)Data acquiring, post-processing and quantification

OCT spectra were acquired at a 100 kHz A-scan rate using customized Labview software. Each B-scan comprised 550 A-scans, resulting in a B-scan rate of 30 Hz that included data acquisition, display and storage. Postprocessing was implemented by customized Matlab™code with standard functions including DC subtraction, dispersion compensation, wavelength-to-k-space interpolation, Hann windowing, and FFT^32,33^. The results were then further processed by averaging or other analysis as indicated. The raw spectrum of each A-scan acquired with OCT and APM-OCT was processed in an exact same way to create images in the spatial domain for comparison.

A metric, normalized speckle contrast (NSC) was used to quantified compare the speckle noise suppression effect between images. It is defined as: the standard deviation (s.d.) of the image intensity in a given region divided by the mean image intensity of the same region. For concise purpose, speckle contrast, instead of its full name, was used in the main text.

The registration of *in vivo* imaging B-scans was done either by ImageJ TurboReg / StackReg plugin^34,35^(for Bscan average), or the phase variance OCT software developed to do intensity averaging and/or blood vessel map extraction^36,37^(for volume data average).

### (3)Wavefront sensorless (WFSL) adaptive optics aberration correction

The image beam at the mouse pupil has a diameter of 0.93μ, a size for which the ocular aberration is nonnegligible. The mouse eye’s aberrations were first corrected using wavefront sensorless (WFSL) aberration correction software^32,38,39^with an image intensity-based searching (SI: Fig. S3). The software automatically calculates the brightness in a user-defined region of interest (ROI) layer, while varying the amplitudes of the DM in Zernike space (ANSI standard^40^) over a search range. After the search process found the optimal mirror configuration for correcting the aberrations of the individual eye, the mirror configuration was loaded into the Labview-based data acquisition software.

### (4)Aperture phase modulation

In an optimized AO system, the DM define a wavefront across the system aperture to corrects aberrations, so as to approach diffraction-limited performance for the system NA, resulting in the most compact point-spread function possible for that NA^27^. The aperture phase distribution was modulated about its optimum AO configuration by random displacements of the mirror segments using a uniform distribution centered on zero, with displacement ranges from 0 (no displacement) to 1.0 μ m (0 ± 0.5 μ m). Histogram analysis of the mirror segments illustrate the uniform distribution of the displacements (Fig. 1(d)). Covariance analysis of the mirror position matrix after 100 trials showed that the mirror segment displacements are uncorrelated (Fig. 1 (a) top right inset).

### (5)3D PSF of the AO-OCT system

In a scanning imaging system, the 3D distribution of power at the focal point in the sample defines the system’s point-spread function. For diffraction-limited systems employing non-coherent light sources and having a circular aperture, the 3D PSF has an analytic form ^41^that can be approximated by a 3D ellipsoid. In OCT, which relies on partially coherent light for interferometry, beam propagation into the sample is governed by the NA of the system in the same manner as for non-coherent light, but the axial direction was further sectioned by the coherence length which is inversely proportional to source bandwidth^33^. In the AO-OCT system used here the PSF has a calculated axial (coherence) length of ~ 2.5 μ m (in tissue, assuming a refraction index of 1.35). In OCT the sampling unit is the A-scan, which provides an axial profile of the backscattering light along the beam propagation axis. While the coherence length of the PSF is invariant with A-scan depth, the lateral (x-, y-) width of the PSF varies according to the NA, being wider away from the center focus. This lateral variation can be particularly notable in AO-OCT, where higher NA is employed, diminishing both the lateral resolution and the power density (imaging brightness) at axial distances away from the center focal plane.

### (6)Timing and Scanning protocol

The configuration of the DM was changed immediately before each B-scan. For AO-OCT, the DM was flatten for resolution target imaging or optimized for aberration-corrected mouse retinal imaging; For APM-AO-OCT, the DM adds random mirror segment displacements on top of the optimal mirror shape for AO-OCT (SI: Fig. S4 (a)-(b)).

1. For the B-scan based comparison between AO-OCT and APM-AO-OCT, the x-scanner repeatedly scanned the same line on the sample. Number of N (N= 100 for *ex vivo* imaging, N=1 for *in vivo* imaging) AO-OCT and APM-AO-OCT were acquired one right after the other one (SI: Fig. S4 (c)).
2. For the enface comparison, number of N (N= 20, 50 or 100 for *ex vivo* imaging, N=50 for *in vivo* imaging) OCT and SM-OCT B-scans were acquired in same location one after one, then the y-scanner move to the next location to repeat the previous process until it covered the ROI (SI: Fig. S4 (d)).

Note that, there is only a N/30 (30Hz B-scan rate) second difference between AO-OCT and APM-AO-OCT data set, to ensure strict comparison.

### (7)Animal Handling

All mice husbandry and handling were in accordance with protocols approved by the University of California Animal Care and Use Committee, which strictly adheres to all NIH guidelines and satisfies the Association for Research in Vision and Ophthalmology guidelines for animal use. Adult pigmented C57BL/6J and albino BALB/c mice were obtained from Jackson Laboratories and maintained on a 12:12, ~100-lux light cycle. During our measurements, mice were anesthetized with the inhalational anesthetic isoflurane (2% in O_2_), and their pupils dilated with medical grade tropicamide and phenylephrine. A contact lens and gel (GelTeal Tears, Alcon, U.S.) was used to maintain the cornea transparency for *in vivo* retinal imaging^42,43^.

### (8)Supplementary movies

Key data were made into tiff stacks first, and then saved as .AVI movie using Image J for demonstration.

### (9)Data availability

The data that support the findings of this study are available from the corresponding author on request.

## Results

### Effect of aperture phase modulation and hypothesized mechanism of speckle noise reduction

As described in the Introduction, speckle noise in OCT images arises from the interference between scattering light from different scatters within the PSF and is observed as voxel-to-voxel intensity fluctuations in the image ^6,44^. In a single OCT B-scan of a Lambertian target (Fig. 2a), the speckle pattern predominates to the extent that no structure can be discerned below the surface. Averaging 100 B-scans with unchanged DM configurations does little to suppress the speckle since the speckle pattern doesn’t change, as dictated by physics, given that the sample and the underlying scatters are immobile for non-biological sample (Fig. 2b).

**Fig. 2.**
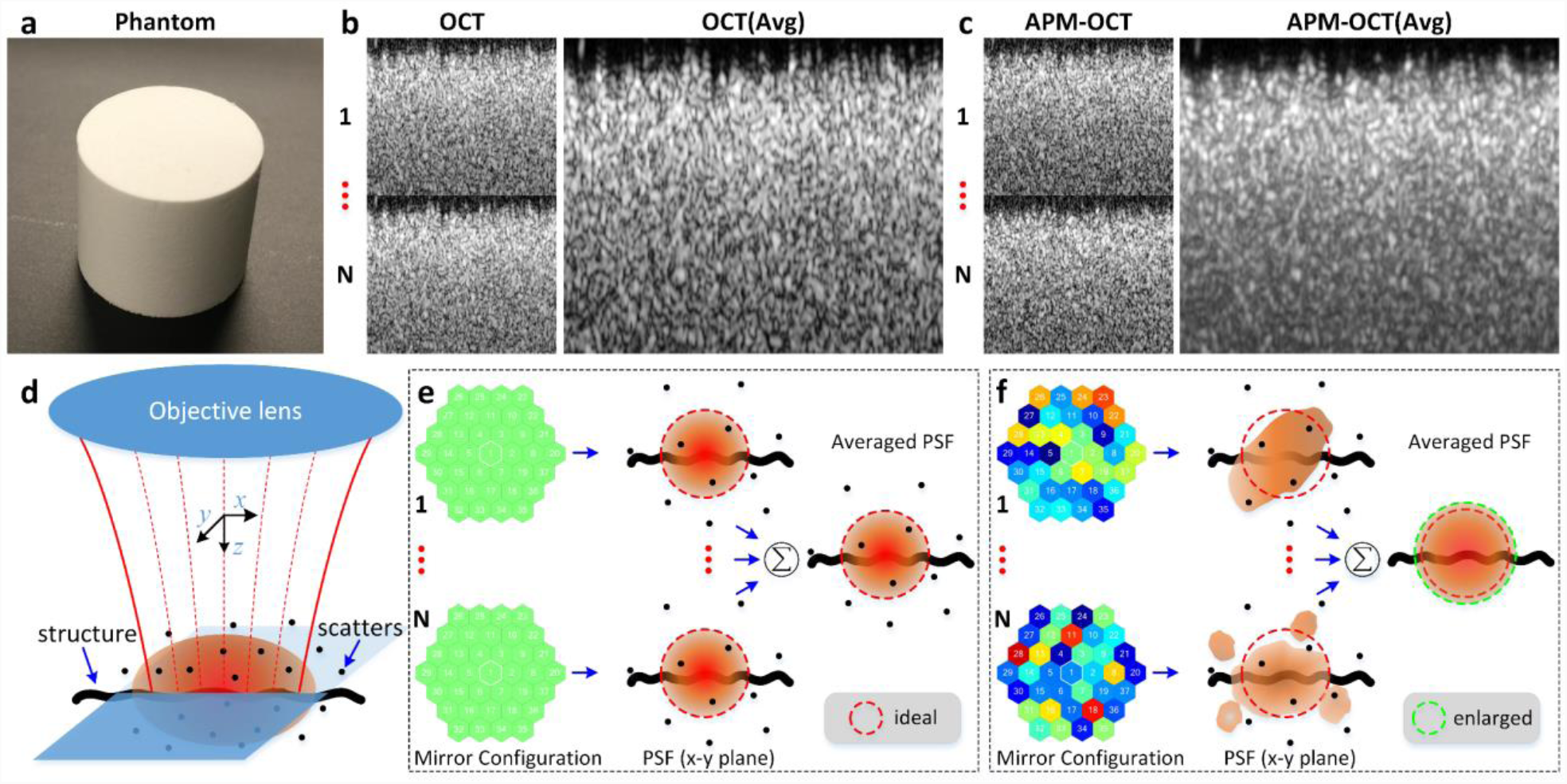
Effect of aperture phase modulation and hypothesized mechanism of speckle noise reduction. **a**, the reflectance standard phantom (Fluorilon 99W, Avian Technologies LLC). **b**, individual (1…N) and 100-frames-averaged OCT B-scans. **c**, individual (1…N) and 100-frames-averaged APM-OCT B-scans. **d**, representation of the in-focus 3D OCT PSF (reddish ellipse). **e**, when mirror was configured as flat mode, a static PSF always selects same scatters set; **g**, when mirror was configured as ‘random’ mode, a dynamic PSF selects different scatters sets. Avg: 100-frames-averaged.

The OCT imaging system has a deformable mirror (DM) whose actuators have a rapid response time, and so afford the possibility of manipulating the wavefront phase at the system aperture. If prior to the collection of each B-scan the DM mirror facets are randomly displaced a sub-micron distance, the speckle pattern changes between B-scans, further averaging will suppress the speckle (Fig. 2c).

For non-living tissue, when the DM of the AO-imaging system is optimized, the PSF realizes its most compact form in the sample (Fig. 2e, PSF, x-y plane) and does not change, so that scatters set sample by the PSF is always same resulting un-changed speckle pattern, explaining why the average B-scan is very similar to any individual scan. Conversely, random displacements of the DM segments from their optimum positions alter the wavefront phase across the aperture, resulting in a PSF that is distorted from the optimum to varying in shapes, intensity distributions and/or extents (Fig. 2f, PSF, x-y plane). This altered PSF will probe a different set of scatters which creates un-correlated speckle pattern between B-scans, while still including a portion of structures (Fig. 2f, thick wavy line, larger than the PSF in either of 3 dimensions) that was sampled by the undistorted PSF. Averaging over a population of B-scans taken with a different DM patterns can thus reduce speckle while preserving signal from the structures (Fig. 2f).

APM-OCT clearly holds promise in reducing speckle noise but faces several challenges. One of these is the inherent conflict between the goal of reducing speckle noise and that of maintaining maximal image resolution. A second challenge is that the potential number of DM configurations is vast: for a mirror with 37 segments and a uniform, only 11-step distribution over the displacement range, the total number of possible configurations is very large (11^37^). A third problem is that it also reduces signal intensity. Practical implementation of APM-OCT as a method of speckle noise reduction must provide an efficient way of selecting a manageable subset of the mirror configurations that also resolves the conflict between speckle noise reduction, and preservation of resolution and signal strength.

### Finding the DM displacement range that both reduces speckle and preserves resolution

To address the conflict between reducing speckle noise and preserving resolution we performed OCT imaging on a printed 1951 USAF resolution test target (Newport, Irvine, CA, U.S.). The B-scan of the target averaged from an ensemble of 100 scans taken with no DM modulation exhibits speckle noise (Fig. 3b), while the average of 100 scans taken with each DM facet displaced randomly over a 0.3 μ m range (0 ± 0.15 μ m) shows strongly reduced speckle (Fig. 3c). The dependence of speckle noise reduction on the number of averaged B-scan and mirror displacement range was quantified by calculating the normalized speckle contrast ^1,26^(Fig. 3d). Here the displacement ranges were varied from 0 (no displacement) to 1.0 μ m with a uniform distribution centered on zero. Speckle contrast rapidly declined with the increased displacement range and/or number of B-scan averaged (Fig. 3d), approaching an asymptotic value. Image resolution loss and speckle contrast reduction from these experiments were then compared as a function of the DM displacement range (Fig. 3e): the curves for the two measures cross at a displacement range of ~ 0.3 μ m, implying that an arrangement of mirror displacements derived from a distribution around 0.3μ m is the best choice for simultaneously preserving resolution and reducing speckle noise for this sample.

**Fig. 3.**
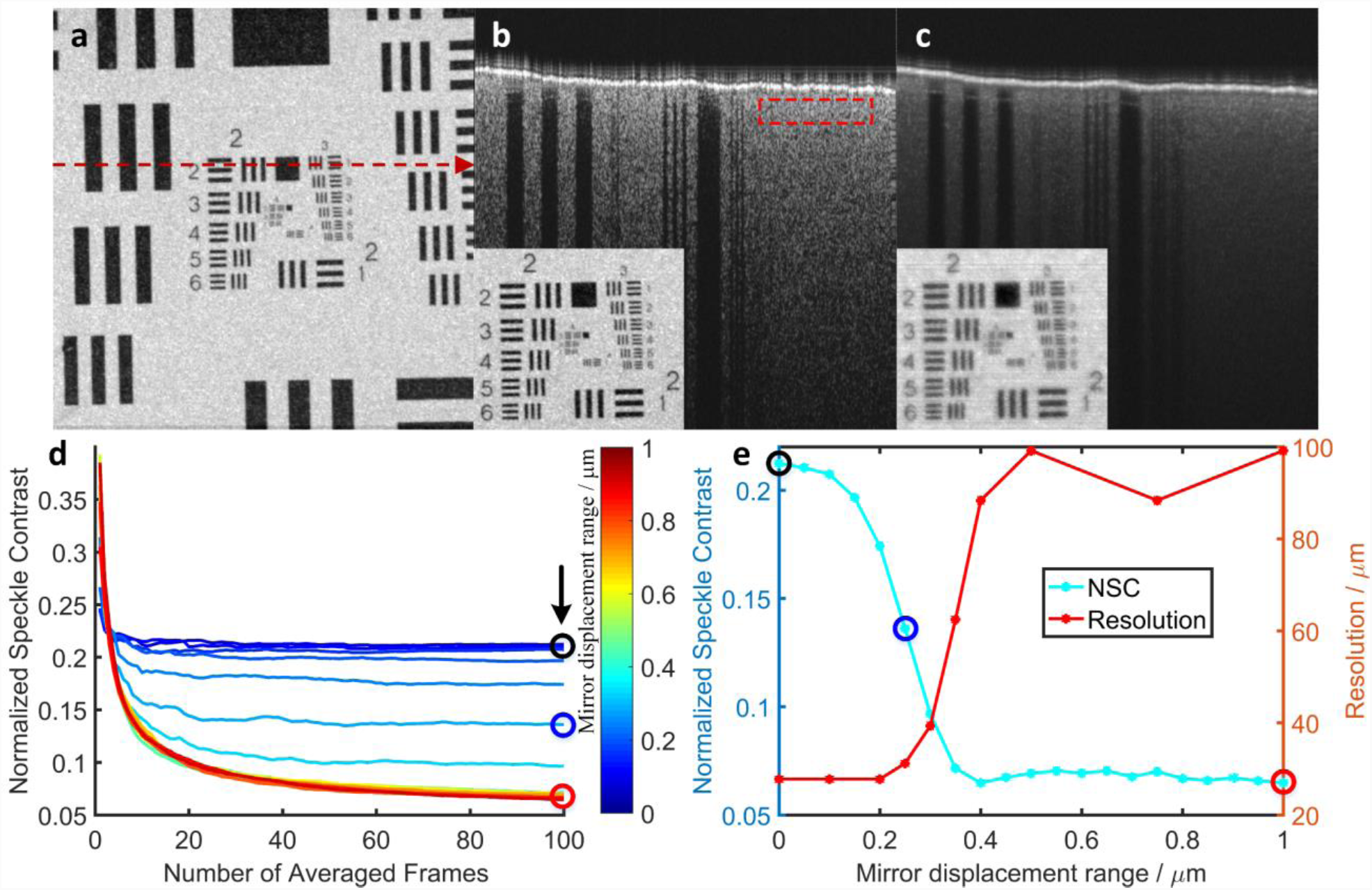
Finding an optimal displacement range for minimizing speckle while preserving target resolution. **a**, OCT enface of a 1951 USAF resolution test target. The red dashed line indicates the OCT Bscans location for panels (b) and (c)). **b**, OCT B-scan obtained by averaging 100-frames when the DM was set to ‘flat’ (red dashed rectangle: region used for speckle contrast quantification; inset: *enface* projection). **c**, APM-OCT B-scan obtained by averaging 100-frames when the DM was random modulated with (**a**) displacement range of 0.3 μm (0 ± 0.15 μm) (inset: *enface* projection) [SI: Video_S2]. Note that, the second layer of the target become visible [arrows pointed in SI: Fig. S5]. **d**, Speckle contrast as a function of the averaged B-scans numbers for different random mirror displacement ranges; the color bar specifies the displacement range [SI: Video_S3]. **e**, Speckle contrast (cyan, same from the arrow pointed data in panel (**d**)) and resolution (red) compared in averaged B-scans as a function of the mirror displacement range [SI: Fig. S6]. For determining resolution only 20 frames were used to save time, since this number reduced speckle contrast by more than 80% by comparing with 100-frames averaged.

### Defining an optimum subset of deformable mirror configurations to preserving the resolution and signal intensity

While the above results show that an optimum mirror displacement range of ~ 0.3 μm can be found (Fig. 3), there is still considerable resolution loss, and substantial signal intensity loss (Fig. 3c, SI: Fig. S6). But the huge number of potential DM configurations makes it is very inefficient to search for a global optimum DM configuration to mitigate the resolution and signal intensity loss. In analyzing the results, we found that the APM-OCT image intensity varied substantially between B-scans quite a lot (over 3 times, linear). Based on general principles, it could be expected that the brighter an individual image, the less distorted was the underlying PSF, suggesting that the subset of mirror displacement patterns yielding the brightest images might correspond to a set of minimally distorted PSFs.

To examine this hypothesis, we generated an ensemble of 1000 B-scans for a mirror displacement range of 0.3 μm and sorted them by their averaged signal intensities (Fig. 4a). Then, sequentially apply the same mirror configurations across the ROI (locations indicated in panel (a) inset). While average intensity varied somewhat for B-scans taken at different positions of the target (Fig. 4b, arbitrary offsets were added for clear display purpose), the overall OCT signal plots were very similar, consistent with the idea that the shape of the plot was dictated by the PSFs corresponding to each mirror configuration, rather than by properties of the sample. The subset of first 100 mirror configurations (Fig. 4 (a) blue region marked), corresponding to top 10% brightest images, were selected for further examination. We compared the ability of the selected top 10% subset of mirror configurations to reduce speckle noise with that of an equal number of random configurations for displacement ranges between 0 and 1.0 μm, and found the selected subset of configurations performed almost as well (Fig. 4c). Remarkably, however, the selected subset of configurations provided resolution up to ~3-fold greater than the randomly generated configurations (Fig. 4d). Even through, Fig. 4 d inset shows there is a continuous contrast loss although the resolution readout doesn’t change. In conclusion, the “top 10%” subset of mirror configurations with displacement range of around 0.3 μm satisfies the triple constraints of greatly reducing speckle noise while simultaneously maximally preserving resolution and signal strength. More generally, the approach provides a rapidly implemented method for programming a deformable mirror to achieve these goals.

**Fig. 4.**
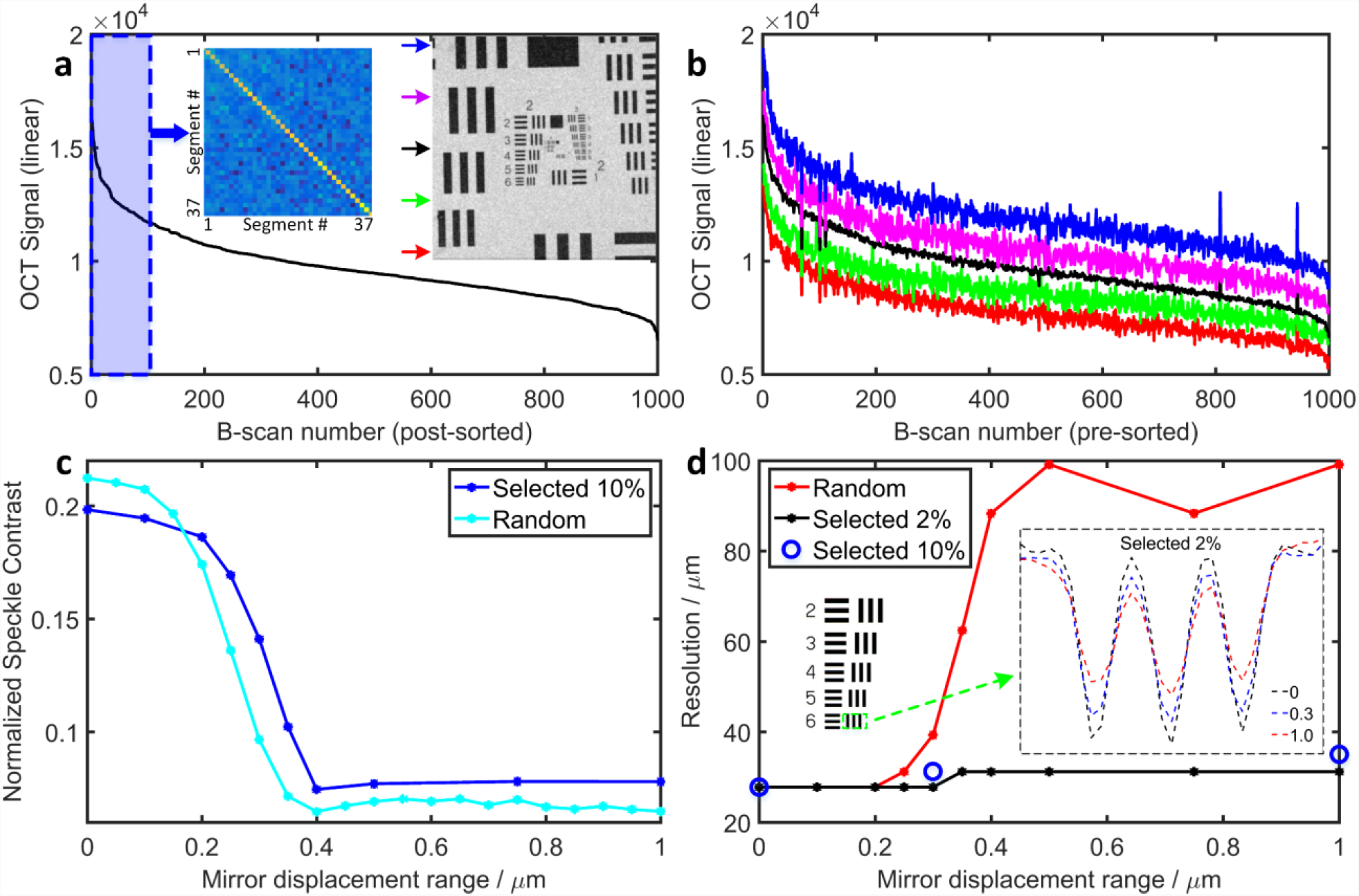
A subset of the mirror configurations reduces speckle while preserving resolution and signal strength. **a**, Average intensity of 1000 APM-OCT B-scans plotted in descending order (the mirror displacement range was 0.3 μm). Left inset: covariance analysis of the top 100 mirror configurations). Right inset: *enface* test target image indicates the B-scan locations for (**b**). **b**, APM-OCT signals from 1000 B-scans with same mirror configuration in (**a**) at different positions, showing a similar form with (**a**). **c**, Speckle contrast comparison for 100-frames-averaged APM-OCT images obtained with random and the selected “top 100” mirror configurations (blue region marked in (**a**)); **d**, Resolution plotted as a function of DM displacement range for different configurations: random (red symbols and line), the top 10% (blue circles), or the top 2% (black symbols and line). The resolutions achieved with the selected mirror configurations is always better than that with random configurations for displacement range greater than 0.25 μm and is asymptotically ~ 3-fold better [SI: Fig. S6]. Left inset: location on target grid for results plotted in right inset. Right inset: vertically averaged cross-section OCT signal changes for different displacement ranges using the selected 2% configurations, showing there is a continuous contrast loss.

### *In vivo* application of APM-AO-OCT reduces speckle efficiently and reveals novel structure

To examine the *in vivo* applicability of APM-AO-OCT we imaged the retinas of Balb/c mice using an interlaced B-scan acquisition protocol in which successive scans were acquired with or without APM (Speckle contrast quantification for in vivo mouse retina was provided in SI: Fig. S7). Single B-scans exhibited substantial speckle that obscured even the highly scattered and extended structures, with little noticeable difference between scans taken with and without APM (Fig. 5a, d). The averages of 32 B-scans with and without APM had noticeably reduced levels of speckle (Fig. 5b, e); notably, extended structures such as the ELM and Bruch’s membrane appeared clearer in the image generated with APM. We quantified the speckle contrast in the region of the B-scans corresponding to the inner plexiform layer (IPL; dashed rectangles in Fig. 5a, d), as this region was bright, but showed no apparent structure. This quantification revealed that the averages of 32 scans taken with APM-AO-OCT had a reliably reduced level of speckle contrast relative to average of 32 scans taken with AO-OCT alone (Fig. 5b, e; Fig. 5g, arrow). The reduction in speckle contrast was evident for all sample sizes between 10 and 1000 (Fig. 5g). The AO-OCT results are consistent with previous observations showing that averaging *per se* leads to reduction in speckle contrast in *in vivo* imaging^24,25^. This reduction was hypothesized to arise from the movements of subcellular organelles whose scattering gives rise to speckle. This hypothesis is supported by our observation that averaging of AO-OCT images of non-living targets (e.g., Fig. 2, 3) does not *per se* much reduce speckle. Nevertheless, APM-AO-OCT more efficiently reduces speckle. Thus, the average of 32 scans with AMP-AO-OCT (Fig. 5e) appears comparable to that of 1000 scans taken with AO-OCT alone (Fig. 5c).

**Fig. 5.**
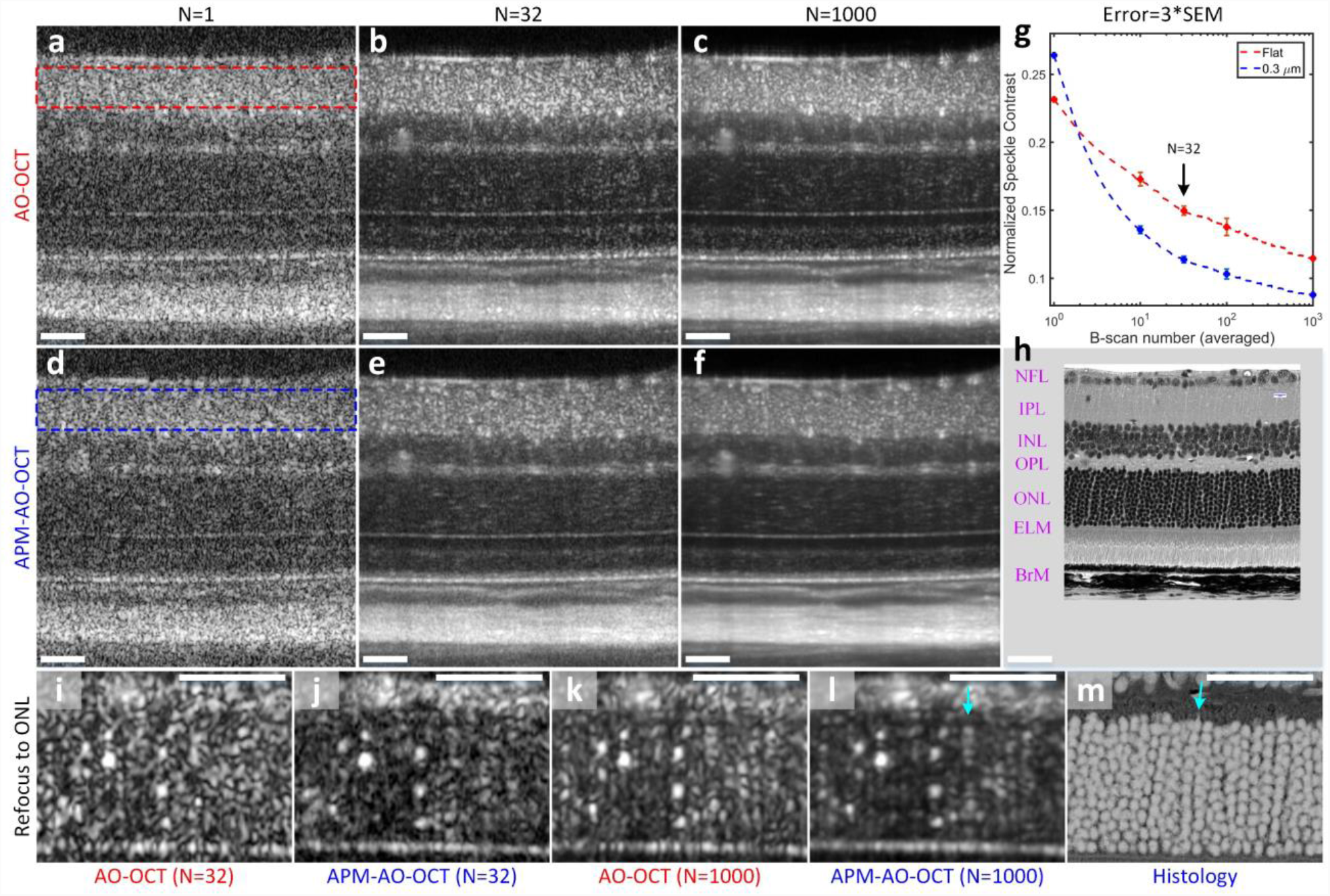
Comparison of the efficiency of the averaging of APM-AO-OCT vs AO-OCT results in reducing speckle and revealing novel cellular structure *in vivo*. **a**-**c**, AO-OCT B-scans with N representing the number of images averaged. **d**-**f**, APM-AO-OCT B-scans with sample averaging corresponding to that used in panels (**a**-**c**). The data in these panels were acquired with interlaced protocol. The focus of the AO-system was set to the IPL. The retinal layers are indicated in (**h**), which is provided at the same scale as the OCT B-scans. **g**, Normalized speckle contrast of the IPL, for AO-OCT (red rectangle in a; red symbols and line in g) and for APM-AO-OCT (blue rectangle in d; blue symbols and line in g), plotted as function of the number of B-scan averaged. **h**, Retinal plastic section of a C57Bl/6 mouse imaged with a 40X objective in a Nikon A1 microscope. **i**-**l**, Averaged B-scans with the focus of the AO system shifted to the ONL; the shifted focus both increases the overall brightness of the images and narrows the width of the ONL scattering spots relative to those in panels (**a**-**f**). **m**, Histology of the ONL from (**h**) presented with inverted contrast and magnified so as to have the same scale as panels (**i**-**l**); scale bar 50μ. Cyan arrow in (**l**) points to a periodic series of spots which is very similar to stacks of rod cell bodies in m. Abbreviations: NFL nerve fiber layer, IPL inner plexiform layer, INL inner nuclear layer, OPL outer plexiform layer, ONL outer nuclear layer, ELM external limiting membrane, BrM Bruch’s membrane. [SI: Video_S4]

In addition to its greater efficiency than pure averaging in reducing speckle noise, APM-AOOCT also serves to increase the confidence with which the experimenter can draw conclusions about structures. To illustrate this point, we compare OCT images taken with the two methods after shifting the focus of the AO-system to the ONL (Fig. 5i-l). The ONL comprises the cell bodies of the photoreceptors, which are developmentally arranged in vertical stacks of 10-11 (Fig. 5h, histology). The average of 32 AO-OCT scans (Fig. 5i) shows spots of increased scattering that might be hypothesized to arise from the photoreceptor nuclei. However, the speckle noise is such that the hypothesis is dubitable. The average of 32 APM-AO-OCT B-scans strengthens the hypothesis (Fig. 5j). The comparison of averages of 1000 B-scans (Figs. 5k, l) leads to even greater conviction that the bright spots arise from rod nuclei: thus, for example, in Fig. 5l one can observe a number of rows of such spots which have the same vertical spacing and in some cases the expected total number as rod nuclei seen in ONL histology (Fig. 5m; contrast-inverted from Fig. 5h). While the hypothesis that photoreceptor nuclei can be visualized with APM-AO-OCT (and to a lesser extent, AO-OCT) needs to be tested further, the evidence from the vertical and lateral spacing as well as size is substantial, and demonstrates the potential for APM-AO-OCT for producing novel discovery. Thus, for example, it is possible that the variation in the brightness of the ONL spots reflects diurnally or otherwise changing structural and/or functional properties of the cell bodies and nuclei.

To explore the full potential of APM-AO-OCT to reduce speckle and uncover structure *in vivo*, we applied the method to volumetric data acquisition, arranging the focus of the AO-system to be at the uppermost retinal layers (Fig. 6a), and comparing AO-OCT with APM-AO-OCT as before. *Enface* presentation of single averaged volume layer showed an enhanced reduction of speckle by APM-AO-OCT (Fig. 6c vs. 6b) and several linear structures (arrows) not discernible in the corresponding AO-OCT image. The averaged single layer about 10μ deeper in the retina (Fig. 6d, e) revealed several regions with intensity greater than the surround which include bright spots. Based on a comparison with published histology (Fig. 6f, Low-power electron microscopy^45^), we hypothesize that these regions represent displaced amacrine cells. Another comparison at the level of the NFL is provided in Fig. 6g, h. Here APM-AO-OCT provides a greater reduction of speckle noise and improved confidence in the discrimination of blood vessels from ganglion cells axon fiber bundles. A potential downside of APM-AO-OCT is that its utility for OCT angiography^36,46^is reduced (Fig. 6i, j). However, this problem can be dealt with the interlaced scanning protocol, as the AO-OCT-alone scans retain the angiographic information (Fig. 6i). Furthermore, the comparison of the averages from the interlaced protocol may lead to insight into the scattering structures seen with the AO-OCT images (compare Fig. 5k and l).

**Fig. 6.**
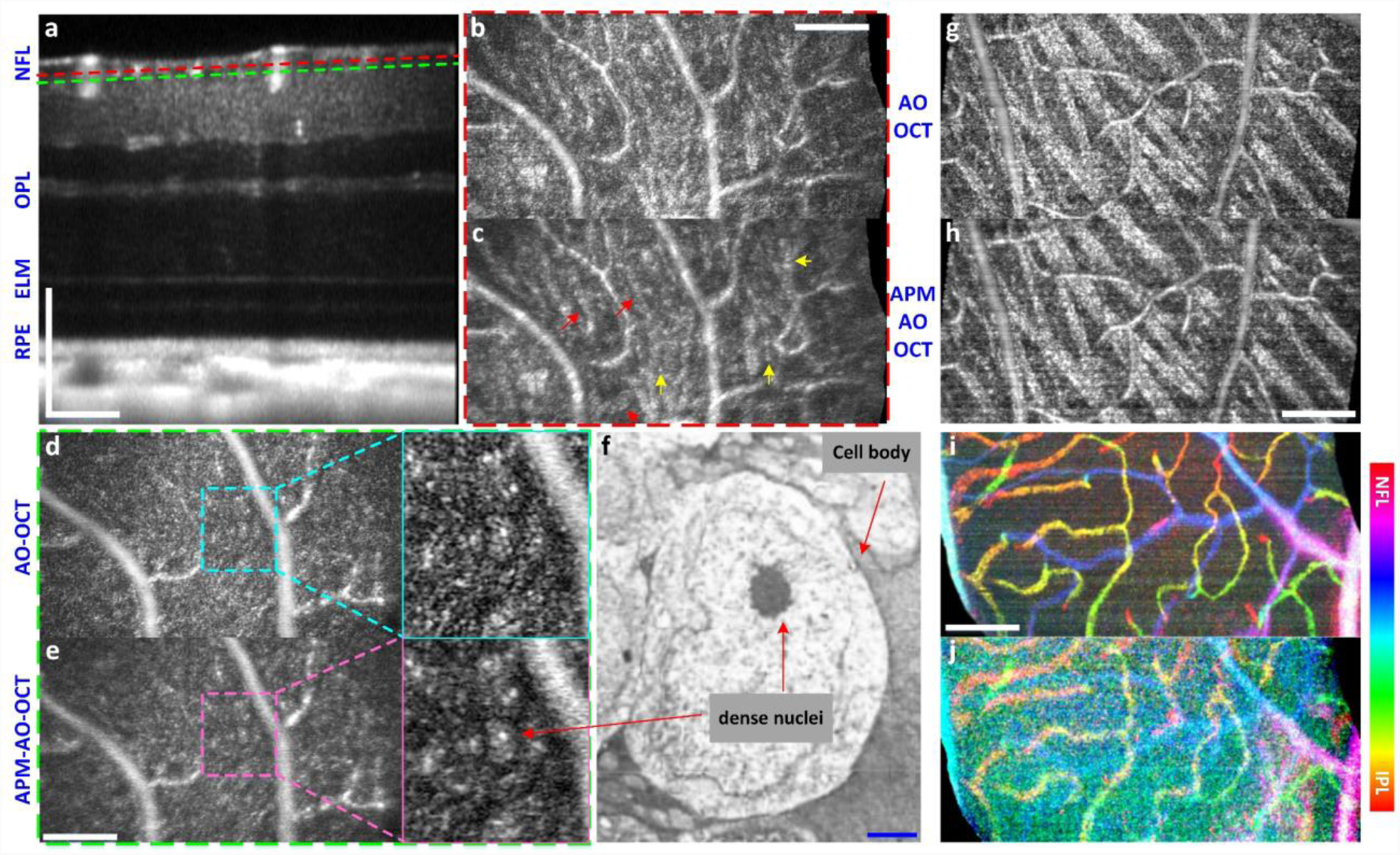
Visualization of cellular scale structures in retinal layers with i*n vivo* volumetric APM-AOOCT. **a,** B-scan from a 560 × 280 × 320μ^3^retinal volume imaged 50 times with interlaced AO-OCT and APM-AO-OCT, aligned and averaged; the AO system was optimized for focus on the outer retina. The dashed lines indicate planes at which enface images were extracted for panels (**b-e**), respectively; **b, c,** *Enface* presentation of a 0.85μ digital section at the depth locus indicated by red dashed line in a for AOOCT (**b**) and APM-AO-OCT (**c**) respectively. Red arrows point to thin line structures that can be excluded as being blood vessels, and likely represent the outermost ganglion cell axons; **d**, **e**, *Enface* presentation of a 0.85μ digital section at the depth locus indicated by green dashed line in a, 10μ deeper into the retina than (**b**, **c**). Magnified presentations reveal relatively brighter (gray) contiguous regions with especially bright dots inclosed; these regions are hypothesized to reveal displaced amacrine cells, which are known to reside in this layer; **f**, Electron microscopic image of an amacrine cell image (from ^45^, with permission). **g**, **h**, *Enface* presentations of 0.85μ digital sections for AO-OCT and APM-AO-OCT with focus on the NFL. Speckle noise reduction by APM-AO-OCT enables more confident discrimination between blood vessels and axon fiber bundles; interlaced protocol; **i**, **j** OCT angiography (phase-variance analysis) with AO-OCT (**i**) and APM-AO-OCT (**j**). The aperture phase modulation substantially reduces the phase- variance signal in the APM-AO-OCT data, while the interlaced AO-OCT data preserves the signal. Scale bar 100μ (white) for all panel except (**f**), where it represents 1μ (blue). Abbreviations: NFL nerve fiber layer, OPL outer plexiform layer, ELM external limiting membrane, RPE retinal pigment epithelium. [SI: Video_S5]

## Summary and Discussion

Adaptive optics has revolutionized image science by enabling image systems to perform at their diffraction limits, and thereby reveal a wealth of novel structure^29^. AO systems operate by actively controlling the wavefront at the system pupil aperture and have been implemented in imaging systems for *in vivo* ophthalmic imaging, including Scanning Laser Ophthalmoscopy (SLO) and OCT systems^47-50^.OCT imaging systems employ partially coherent light sources to extract depth scattering profiles of tissue, and as with all systems that use such sources, are subject to speckle noise, which substantially reduces their signal-to-noise ratio. Here we have presented a novel approach to speckle noise reduction in OCT. This approach exploits small scan-to-scan modulations of the phase at the aperture of an AO-OCT system produced by submicrometer displacements of the segments of a deformable mirror (DM) (Fig. 1). We established that an optimum mirror displacement range can be found which simultaneously greatly reduces speckle noise and maintains image resolution (Fig. 3), and that a small subset of the mirror configurations can further improve resolution and preserve signal intensity (Fig. 4). Finally, we demonstrated APM-AO-OCT can be used *in vivo* to efficiently reduce speckle noise and discover novel structure (Fig. 5, 6).

### Mechanism of APM-AO-OCT: selected perturbations of the system point-spread function

In an OCT system, the point-spread function (PSF) is defined axially by the source coherence length and determined laterally by the NA of the system (Methods). Because the sampling unit in OCT is the A-scan, the lateral extent of the PSF varies with depth, achieving its diffraction-limited minimum at the focal depth. Aperture phase modulation (APM) necessarily perturbs the OCT PSF shape, but primarily affects its *x*-, *y*distribution. The effects of APM on the PSF can be visualized by focusing the OCT beam onto a CMOS camera (Fig. 7). Each of a series of 1000 APM-AO-OCT PSFs exhibit a central power density with random extensions of lower power (Fig. 7a), while a similar sample of 1000 AO-OCT PSFs are identical (Fig. 7c). The “top 100” with brightest maximum intensity of the APM-AO-OCT sample are more compact (Fig. 7b), as further emphasized by comparison of the averages (Fig. 7d, c), and comparison of line scans through the averaged PSF centers (Fig. 7f). This analysis provides support for the hypothesis (Fig. 2) that the averaging of scans taken with APM-AO-OCT efficiently reduces speckle contrast because the randomly distorted PSFs encompass different sets and numbers of scatterers, while the maintained centroid of the PSFs captures information from larger scale structural elements in the sample.

**Fig. 7.**
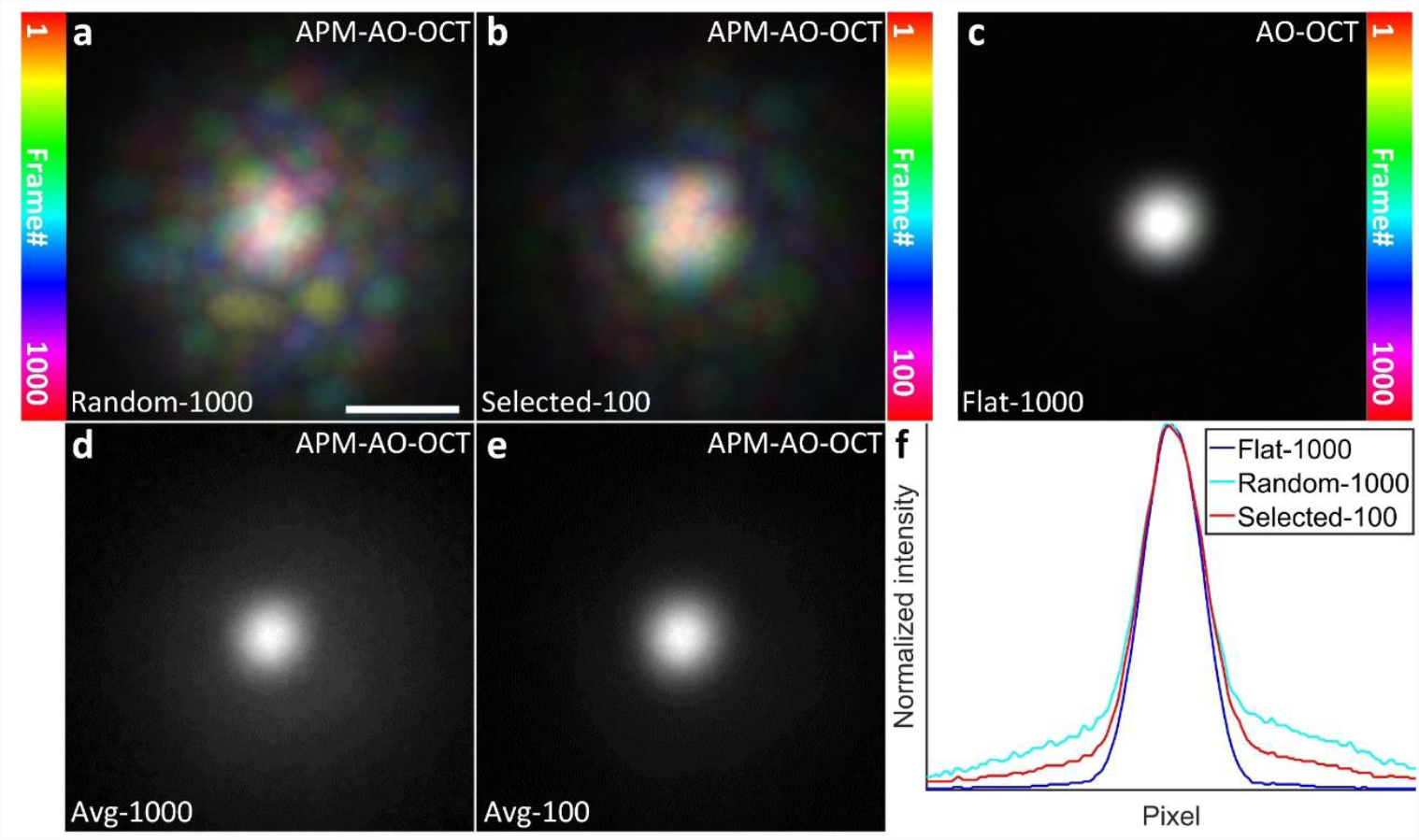
Comparison of the lateral extent of AO-OCT and APM-AO-OCT PSFs at the focus. All PSF images were obtained by focusing the beam onto a CMOS camera. **a**, Color-coded projection of 1000 APM-AO-OCT PSFs produced by mirror segment displacement range of 0.3μ. Colors were assigned according to the position in the series as indicated in the colorbar at left. **b**, Color-coded projection of “top 100” PSFs from the sample of 1000 presented in (**a**). (see also SI Video_S9). **c**, Color-coded projection of 1000 AO-OCT PSFs (no DM modulation); the 1000 PSFs were indistinguishable from one another. **d**, Average of the 1000 APM-AO-PSFs presented in (**a**). **e**, Average of the “top 100” APM-AO-OCT presented in (**b**). **f**, Line profiles of the averaged PSFs; see legend. Note that, these images represent the “1-way” or incoming PSF of the system, whereas in application the effective PSF results from two passes through the system aperture. [SI: Video_S6]

We believe that further insight into the class of mirror displacement configurations that minimize speckle contrast while maintaining resolution and image brightness will be obtained both by additional characterization of the mirror configurations that give optimum performance, and by theoretical analysis of the corresponding perturbed wavefronts. Thus, for example, histogram analysis of the DM segment displacements of the “top 100” configurations as a function of distance from the DM pupil center revealed that the outermost actuators underwent uniform variation over the full range of deformation, while the inner actuator displacements followed a Gaussian distribution with a restricted range. This observation suggests that configurations characterized by the subset of Zernike aberrations of the class *Z*_*j*_^±*j*^ may be especially important in optimizing APM-AO-OCT.

### Comparison with similar methods of speckle reduction

Many different approaches to reducing speckle noise in imaging systems employing coherent light have been proposed ^3,13^(cf. Introduction). Methods such as that of Liba et al. ^2,26^which vary the properties of the incoming beam suffer from lack of control: first, because the distorted PSF is not readily obtained, losing the capability for optimizing; second, because the class of permitted distortions is limited and cannot be easily and precisely varied as would be needed in application with different wavelength and/or samples. APM-AO-OCT overcomes these deficits and gives the experimenter precision control on a very rapid trial-by-trial basis, providing a quantified and repeatable way to further explore and optimize the method; furthermore, it is intrinsically compatible with adaptive optics, offering a natural way to reduce speckle while preserving AO enhanced lateral resolution.

Recently, it was also found that averaging of multiple, precisely aligned volumes for *in vivo* OCT imaging could reduce speckle noise and reveal novel cellular scale structure^24,25^. It was hypothesized that such averaging is effective because of the random movement of sub-PSF size scattering elements in cells. This approach is passive, however, and limited by the time scale and extent of the underlying scatterer motions, which impose a characteristic time window for the decorrelation of the speckle pattern between images. The *in vivo* results presented here confirm the effectiveness of pure averaging, but also show that the active approach of APM-AO-OCT can be considerably more efficient (Fig. 5, 6).

### Future directions

The broad adoption of adaptive optics continues to revolutionize imaging science and has spurred the development of wavefront correctors with increasing numbers of segments and speed, at ever lower cost. In principle, APM-AO-OCT could also be implemented with spatial light modulators (SLM)^51^, digital micro-mirror devices (DMD) ^52,53^, or other deformable mirrors (e.g. AlpAO, BMC Inc., etc.) ^54-56^Exploration of alternative PSF shaping methods^57^and more efficient ways of generating appropriate wavefront deformations should speed the routine implementation of APM-AO-OCT (SI: Fig. S8). Moreover, OCT systems capable of megahertz A-scan rates have been developed ^58,59^. The marriage of modern AO and ultrahigh speed OCT scan technologies should enable routine implementation of APM-AO-OCT in clinical and research settings, enhancing cellular resolution clinical diagnosis as well as basic science discovery. For example, implementation of APM-AO-OCT should lead to broader application of OCT to brain structural imaging^60,61^where the presence of speckle in images remains a major obstacle. Finally, thanks to the improvement in the visualization of subcellular structure, APMAO-OCT should enable a more precise localization and quantification of retinal optophysiological signals, which provide non-invasive, label-free measurement of photoreceptor function^62-64^

## Acknowledgements

Authors would like to acknowledge their funding sources: UC Davis RISE funding, NSF I/UCRC CBSS Grant, NIH grants EY026556, EY02660 and EY012576 (NEI Core Grant), Vision Science T32 EY015387. We thank Prof. Vivek Jay Srinivasan for sharing the OCT processing dll; We thank Prof. Marie Burns and Prof. John S. Werner for help and support. We thank Avian Technologies LLC for providing the Fluorilon 99W reflectance standard phantom used in the experiment.

## Author contributions

P.Z. conceived and designed the research, performed experiments and analyzed data; S.M. contributed system design, building and discussion; E.M. contributed post-processing tools; R.K.M contributed discussion; Y.J and M.V.S. contributed the GPU based real-time OCT processing and WFSL aberration correction software; E.N.P. and R.J.Z. contributed expert advice; P.Z., E.N.P. and R.J.Z. co-wrote the paper.

